# Functional activation of dorsal striatum astrocytes improves movement deficits in hemi-parkinsonian mice

**DOI:** 10.1101/2024.04.02.587694

**Authors:** Wesley R. Evans, Sindhuja S. Baskar, Castro E Costa Ana Raquel, Sanya Ravoori, Abimbola Arigbe, Rafiq Huda

## Abstract

Parkinson’s disease (PD) is characterized by the degeneration of dopaminergic nigrostriatal inputs, which causes striatal network dysfunction and leads to pronounced motor deficits. Recent evidence highlights astrocytes as a potential local source of striatal network modulation. However, it remains unknown how dopamine loss affects striatal astrocyte activity and whether astrocyte activity regulates behavioral deficits in PD. We addressed these questions by performing astrocyte-specific calcium recordings and manipulations using *in vivo* fiber photometry and chemogenetics. We find that locomotion elicits astrocyte calcium activity over a slower timescale than neurons. Unilateral dopamine depletion reduced locomotion-related astrocyte responses. Chemogenetic activation facilitated astrocyte activity, and improved asymmetrical motor deficits and open field exploratory behavior in dopamine lesioned mice. Together, our results establish a novel role for functional striatal astrocyte signaling in modulating motor function in PD and highlight non-neuronal targets for potential PD therapeutics.

## Introduction

Parkinson’s disease (PD) is a progressive neurodegenerative disorder characterized primarily by loss of dopaminergic neurons in the substantia nigra pars compacta (SNpc)^1^. The ensuing loss of dopamine inputs to the dorsal striatum dysregulates local neuronal circuits and leads to the motor symptoms of PD^2–4^. The typical clinical treatment for PD is the dopamine precursor levodopa (L-DOPA) which replaces lost dopamine (DA) to alleviate PD symptoms^5^. However, prolonged treatment with L-DOPA is associated with both resistance to treatment^6^ and dyskinesia^2,5^. Complementary pathways to PD treatment represent possible avenues to improve patient quality of life and outcomes.

Astrocytes are a major subtype of non-neuronal cells that populate all regions of the brain^7^ and modulate diverse neuronal functions including neurotransmitter clearance, ionic homeostasis, cerebral blood flow, and synaptic development and plasticity^8–19^. These glia cells are electrically quiescent but show robust intracellular calcium dynamics^20–23^. Emerging evidence suggests that astrocyte calcium signals reflect local network activity and neuromodulatory inputs^24^. For example, dopaminergic inputs increase astrocyte calcium activity in the nucleus accumbens^25^. In turn, astrocyte calcium activity shapes responses of local neuronal networks^26^. Moreover, chemogenetic modulation of astrocyte signaling in multiple brain regions modulates behavioral output^27,28^, including in the dorsal striatum^29^.

Recent evidence supports the targeting of astrocytes for the treatment of diverse brain disorders^30,31^, including PD^32,33^. Locomotion induces robust calcium activity in Bergman glia of the cerebellum and cortical astrocytes^34^. These responses are mediated by noradrenergic inputs which are absent in the striatum^35–37^. The functional role of astrocytes in movement and the effects of DA loss on striatal astrocyte calcium activity is unknown. Here, we investigated movement-related astrocyte activity using *in vivo* calcium recordings with fiber photometry. We found that striatal astrocytes are robustly activated during locomotion. Further, although astrocyte calcium signals are only moderately disrupted by acute pharmacological blockade of dopamine receptors, unilateral 6-hydroxydopamine (6-OHDA) lesioning of midbrain DA neurons leads to a pronounced deficit. Finally, we found that chemogenetic activation of striatal astrocytes improves motor function in dopamine lesioned mice, demonstrating the potential of leveraging astrocyte mechanisms for treatment of motor symptoms of PD.

## Results

### Dorsal striatum astrocyte calcium activity tracks locomotion

We virally expressed in the dorsal striatum the membrane bound, genetically encoded calcium indicator GCaMP6f-lck under the control of an astrocyte-specific promoter and measured bulk astrocyte calcium activity using fiber photometry (**Fig. 1a**). Post-hoc immunohistochemistry showed extensive overlap between GCaMP6f-lck and the astrocyte marker S100β (**Fig. 1b**). To determine movement-related astrocyte activity, we trained head-fixed mice to voluntarily run on a low-resistance circular disc (**Fig. 1c**). Bulk astrocyte calcium activity closely followed locomotion (**Fig. 1d**). We detected individual locomotion events and aligned astrocyte activity (ΔF/F) to the onset and offset time of locomotion bouts (**Figs. 1d** and **1e**). Activity increased slowly after movement initiation, remained elevated for the duration of movement, and subsided at locomotion offset with a seconds-long time course (**Figs. 1e** and **1f**). Normalized cross-correlation analysis between onset aligned wheel speed and ΔF/F showed that peak astrocyte activity lagged locomotion onset by ∼2s (**Fig. 1g**). Hence, locomotion elicits astrocyte calcium activity with a slow time course.

**Figure 1.**
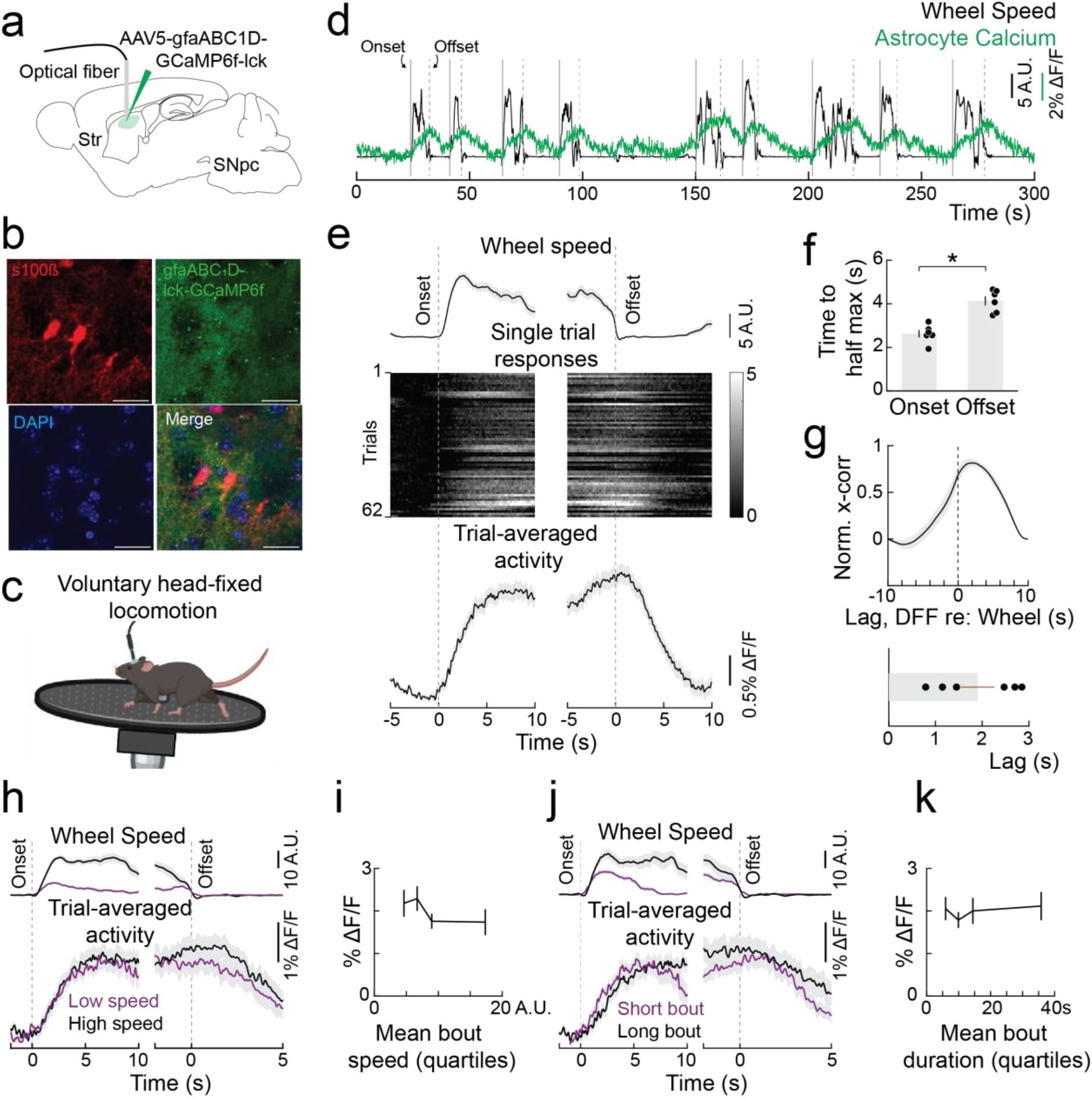
Voluntary locomotion recruits striatal astrocyte calcium signals. (**A**) Experimental schematic showing injection of viral constructs (gfaABC1D-lck-GCaMP6f) and placement of recording fiber optics into the dorsal striatum. (**B**) Confocal image of astrocytes expressing gfaABC1D-lck-GCaMP6f and immunostained for s100β. (**C**) Schematic of fiber photometry recordings during voluntary head-fixed locomotion. (**D**) Astrocyte activity (green) and wheel speed (black) from an example voluntary locomotion session. Dashed lines denote onset and offset times for detected movement bouts. (**E**) Trial-averaged wheel speed (top) and astrocyte activity (middle and bottom) aligned to onset and offset of locomotion bouts (n = 62 trials from 6 mice). (**F**) Comparison of time to half maximum response of astrocyte activity during onset and offset of locomotion bout (n = 6 mice; *p* = 0.028, *z* = -2.20; Wilcon signed-rank test). (**G**) Top, normalized cross-correlation between wheel speed and astrocyte calcium activity. Bottom, average lag between wheel speed and astrocyte activity for individual animals (n = 6 mice). (**H**) Locomotion bouts across all recorded mice were combined and split into quartiles based on average speed between bout onset and offset. Wheel speed (top) and astrocyte activity (bottom) is shown for 1^st^ (low speed) and 4^th^ (high speed) quartiles aligned to bout onset and offset. (**I**) Astrocyte activity measured at the time of locomotion bout offset plotted against mean speed of the 4 quartiles (n = 62 trials from 6 mice; *F*(3,58) = [1.07], *p* = 0.37; one-way ANOVA). (**J**) Same as H, except locomotion bouts were split based on duration of locomotion bouts calculated from onset to offset. (**K**) Same as I, except activity is plotted against the mean duration of the 4 quartiles (n = 62 trials from 6 mice; *F*(3,58) = [0.25], *p* = 0.86; one-way ANOVA).

Next, we determined how astrocyte activity relates to the duration and speed of voluntary locomotion bouts. We split trial-aligned astrocyte activity into quartiles based on the speed of locomotion bouts (**Fig. 1h**). Comparing astrocyte responses for low and high speed (quartiles 1 and 4) locomotion bouts showed that activity reaches a similar peak value under both conditions. Accordingly, there was no significant relationship between speed and ΔF/F measured at the time of locomotion offset (**Fig. 1i**). Similarly, astrocyte activity was not dependent on the duration of locomotion bouts, showing similar responses for short and long bouts (**Fig. 1j** and **1k**). Therefore, movement responses of dorsal striatum astrocytes are not modulated by specific parameters of voluntary locomotion bouts.

### Comparison of locomotion responses between striatal subregions and cell types

We performed fiber photometry recordings in the dorsomedial (DMS) and dorsolateral (DLS) subdivisions of the striatum from neurons or astrocytes to compare movement responses between cell types and striatal subregions (**Fig. 2a**). We trained mice to run on a motorized wheel to control interindividual differences in movement bout parameters (speed and duration) and facilitate comparison of locomotion responses. Locomotion evoked robust calcium increases in both striatal neurons and astrocytes. Across cell types, overall ΔF/F amplitudes did not significantly differ, nor did they differ across DMS and DLS astrocytes (**Fig. 2f**). However, there was a difference in the temporal properties of locomotion responses between neurons and astrocytes. Neuronal calcium activity reached peak values significantly faster than either DMS or DLS astrocytes (**Fig. 2g**). Similarly, offset time after termination of the locomotion bout was significantly faster for neurons than DMS or DLS astrocytes (**Fig. 2h**). There were no significant differences in the temporal properties of DMS and DLS astrocyte responses. Moreover, locomotion-related astrocyte activity did not vary as a function of motorized wheel speed (**Fig. S1**), like that observed with voluntary locomotion (**Fig. 1h** and **1i**). Together, these results show that striatal neurons respond faster during locomotion while DMS and DLS astrocytes show similar locomotion-related activity under these conditions. Given the similarity in response profiles and the importance of DLS in movement control, we performed subsequent fiber photometry experiments on DLS astrocytes.

**Figure 2.**
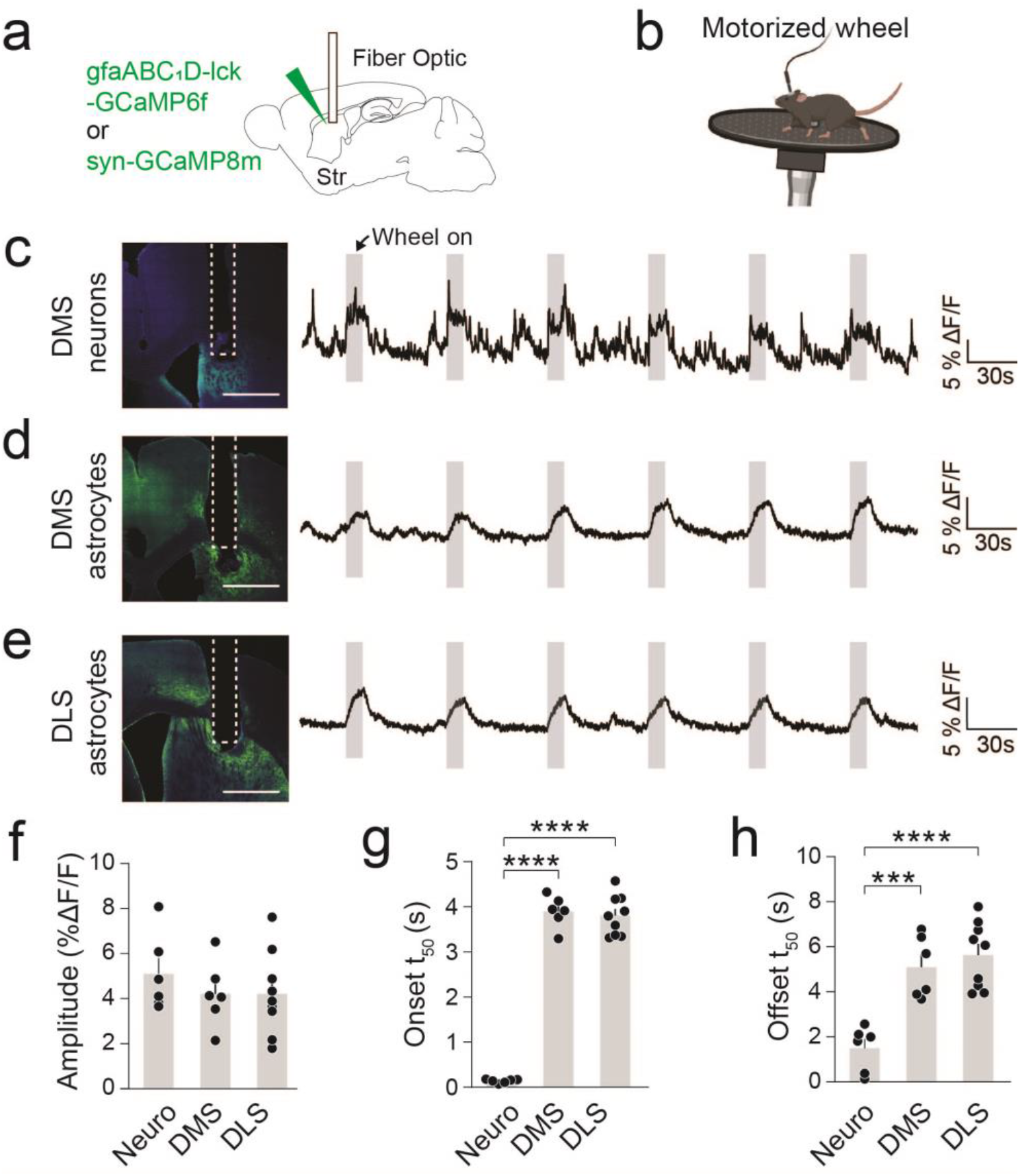
Locomotion recruits dorsal striatal astrocyte activity at a slower timescale than striatal neuronal activity. (**A**) GCaMP was expressed in neurons (syn-GCaMP8m) or astrocytes (gfaABC_1_D-lck-GCaMP6f) and fiber optic cannula was implanted in the dorsal striatum. (**B**) Fiber photometry signals were recorded from head-fixed mice during motorized wheel locomotion. (**C, D, E**) Representative images showing GCaMP expression and fiber placement along with calcium activity during locomotion for dorsomedial striatum (DMS) neurons (**C**), DMS astrocytes (**D**), and dorsolateral striatum (DLS) astrocytes (**E**). Shaded portions show locomotion bouts. (**F**) Comparison of locomotion response amplitude between DMS neurons, DMS astrocytes, and DLS astrocytes (n = 6 mice for DMS neurons, 6 for DMS astrocytes, 9 for DLS astrocytes; *F*(2, 18) = [0.1818], *p* = 0.8353; one-way ANOVA). (**G**) Time to 50% of maximum locomotion response (n = 6 for DMS neurons mice, 6 for DMS astrocytes, 9 for DLS astrocytes; *F*(2, 18) = [4.849], *p* = 1*10^-13^; one-way ANOVA; Sidak’s multiple comparisons test: neurons vs DMS astrocytes *p* = 9.7*10^-13^, neurons vs DLS astrocytes *p* = 3.1*10^-13^, DMS astrocytes vs DLS astrocytes *p* = 0.95). (**H**) Time to 50% offset after end of locomotion (n = 6 for DMS neurons mice, 6 for DMS astrocytes, 9 for DLS astrocytes; *F*(2, 18) = [19.00], *p* = 3.7*10^-5^; one-way ANOVA; Sidak’s multiple comparisons test: neurons vs DMS astrocytes *p* = 5.4*10^-4^, neurons vs DLS astrocytes *p* = 4.0*10^-5^, DMS astrocytes vs DLS astrocytes *p* = 0.83).

### Effect of dopamine loss and L-DOPA on astrocyte locomotion responses in hemi-parkinsonian mice

Loss of dopamine inputs to the dorsal striatum contributes significantly to the motor phenotypes of PD. We determined the role of dopamine signaling in locomotion-related astrocyte responses. Mice were given an I.P. injection of saline as a control or the broad-spectrum DA receptor antagonist flupenthixol (5mg/kg) before locomotion on the motorized wheel (**Fig. S2a**). We performed these experiments at variable duration of wheel movements to broadly match bout durations observed during spontaneous head-fixed locomotion (2, 5, 10, and 20s). Flupenthixol moderately reduced the amplitude of locomotion responses at all tested wheel running durations (**Figs. S2b-d**), reduced the onset time of the response for the 20s bout condition (**Fig. S2e**), and had no effect on the response offset time (**Fig. S2f**). Hence, acute inhibition of dopaminergic signaling reduces astrocyte activity.

To better understand how astrocyte activity is affected by dopamine loss, we employed the widely used 6-OHDA hemi-parkinsonian model^35^. Mice were stereotaxically injected with the neurotoxin 6-OHDA (lesioned) or vehicle control (sham) into the medial forebrain bundle (MFB) to lesion dopaminergic neurons in the left hemisphere (**Fig. 3a**). Post-hoc immunohistochemistry against tyrosine hydroxylase (TH) showed a significant loss of SNpc dopamine neurons and striatal dopaminergic innervation on the lesioned side as compared to the un-injected side (**Figs. 3b-d**). We recorded astrocyte locomotion responses in the dorsal striatum 4-5 weeks after dopamine lesioning. Compared to sham mice, there was a pronounced decrease in astrocyte activity across all but the shortest (2s) tested running durations (**Figs. 3e-g**). The temporal profile of astrocyte responses was not affected for short locomotion durations (2-10s). However, dopamine loss significantly reduced the time to reach 50% of the peak response for the 20s locomotion condition (**Fig. 3h**). 6-OHDA lesioning produced a higher reduction in astrocyte locomotion activity compared to acute dopamine receptor antagonism with flupenthixol (6-OHDA normalized to sham: 42.06±6.1% (n = 9); flupenthixol normalized to saline: 82.8±11.1%; p = 0.01 (n = 9), Wilcoxon ranked-sum test). This suggests that the reduction in astrocyte activity in lesioned mice reflects not only loss of dopamine signaling but possibly also substantial remodeling of the striatal circuit commonly observed with dopamine loss^36^ and other factors.

**Figure 3.**
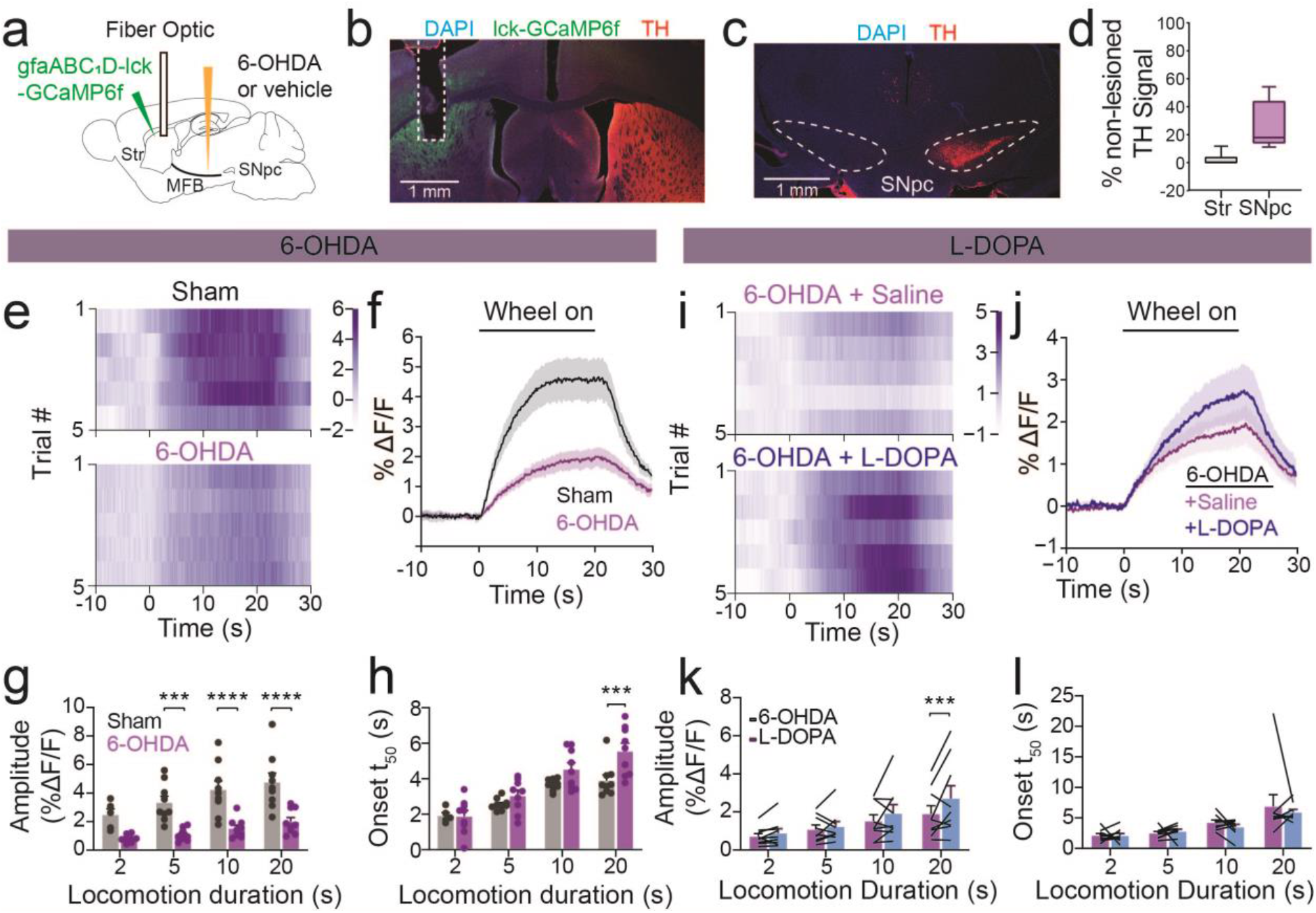
Dopamine loss broadly reduces locomotion-related striatal astrocyte calcium, which is partially improved with L-DOPA treatment. (**A**) Experimental setup showing unilateral viral and 6-OHDA injection sites and recording fiber placement. (**B**) Representative image showing loss of striatal TH immunofluorescence, expression of gfaABC1D-GCaMP6f-lck, and fiber optic implant site on the lesioned side. (**C**) Loss of TH immunoreactivity in the SNpc on the lesioned side. (**D**) TH immunofluorescence in the striatum and SNpc of the lesioned side relative to the non-lesioned side. (**E**) Colormap plot showing single trial responses for 20s locomotion bouts in example sham (top) and dopamine lesioned (bottom) animals. (**F**) Trial-averaged locomotion responses for the 20s duration condition across all recorded mice (n = 9 sham and 9 lesioned animals). (**G**) Locomotion response amplitude across the tested durations for sham and dopamine lesioned animals (n = 9 mice per group; main effect of duration *F*(3, 44) = [51.59], *p* = 2*10^-14^; main effect of drug treatment *F*(1, 16) = [16.33], *p* = 9.5*10^-4^; interaction *F*(3, 44) = [9.514], *p* = 6.0*10^-5^: Sidak’s multiple comparisons test: 2s *p* = 0.29, 5s *p* = 9.6*10^-4^, 10s *p* = 5.5*10^-5^, 20s *p* = 4.4*10^-5^, two-way mixed-effects ANOVA). (**H**) Time to 50% of peak locomotion was reduced by dopamine lesioning during longer locomotion bouts (n = 9 mice per group; main effect of duration *F*(3, 44) = [54.91], *p* = 6.0*10^-15^; main effect of treatment *F*(1, 16) = [5.474], *p* = 0.033; interaction *F*(3, 44) = [4.200], *p* = 0.0107: Sidak’s multiple comparisons test: 2s *p* = 0.99, 5s *p* =0.67, 10s *p* = 0.16, 20s *p* = 8.4*10^-4^, two-way mixed-effects ANOVA). (**I**) Single trial responses from a 6-OHDA lesioned animal injected with either saline or L-DOPA/Benserazide (10mg/kg, 12mg/kg) (bottom). (**J**) Mean locomotion responses for saline (black) and L-DOPA (purple) conditions (n = 9 mice). (**K**) Response amplitude in lesioned animals with L-DOPA and saline injections (n = 9 mice; main effect of duration *F*(3, 24) = [16.24], *p* = 5.6*10^-6^; main effect of treatment *F*(1, 8) = [2.838], *p* = 0.13; interaction *F*(3, 24) = [3.355], *p* = 0.036: Sidak’s multiple comparisons test: 2s *p* = 0.79, 5s *p* =0.88, 10s *p* = 0.11, 20s *p* = 2.8*10^-4^, two-way repeated measures ANOVA). (**L**) Effect of L-DOPA on time to 50% of peak response (n = 360 trials from 9 mice; main effect of duration *F*(3, 24) = [10.85], *p* = 1.1*10^-^; main effect of treatment *F*(1, 8) = [0.2747], *p* = 0.61; interaction *F*(3, 24) = [3.355], *p* = 0.72: Sidak’s multiple comparisons test: 2s *p* = 0.99, 5s *p* =0.99, 10s *p* = 0.87, 20s *p* = 0.73, two-way repeated measures ANOVA).

L-DOPA is used as a dopamine replacement therapy in PD. We tested how L-DOPA treatment in dopamine lesioned animals affects astrocyte locomotion responses. L-DOPA (10 mg/kg, I.P.) moderately increased astrocyte activity during the 20s locomotion condition as compared to saline administration (**Figs. 3i-k**). However, there was no effect for shorter duration bouts (**Fig. 3k**). We also did not detect an effect of L-DOPA on the temporal profile of astrocyte responses (**Fig. 3l**).

Together these results suggest that while dopamine signaling contributes only moderately to functional astrocyte responses under basal conditions, dopamine loss is associated with pronounced reductions in astrocyte calcium activity.

### Chemogenetic activation facilitates astrocyte calcium activity

Astrocytes express a wide array of G-protein coupled receptors (GPCRs) that couple to diverse intracellular second messenger systems and modulate intracellular calcium activity^37–41^. We tested the effect of engineered GPCRs^42^ on astrocyte activity and parkinsonian motor phenotypes. We virally expressed in each striatal hemisphere either the Gi DREADD actuator GFAP-hM4Di-mCherry or GFAP-mCherry as fluorophore control (**Fig. 4a**). Cohorts of mice were given I.P. injections of saline or the DREADD agonist clozapine-N-oxide (CNO; 3mg/kg) and brains were harvested 2 hours after injections. Post-hoc immunohistochemistry showed that DREADD expression was predominantly colocalized with the astrocyte-specific marker S100β but not the neuronal marker NeuN (**Fig. S3a** and **S3b**). In CNO treated mice, most DREADD-expressing astrocytes were co-labeled with c-Fos, a marker for intracellular calcium activity^29^ (**Fig. 4a**, **b**). However, astrocytes expressing only mCherry did not show c-Fos labeling. In mice treated with saline, little c-Fos expression was observed in DREADD or mCherry expressing astrocytes (**Figs. 4b and S3c-e**).

**Figure 4.**
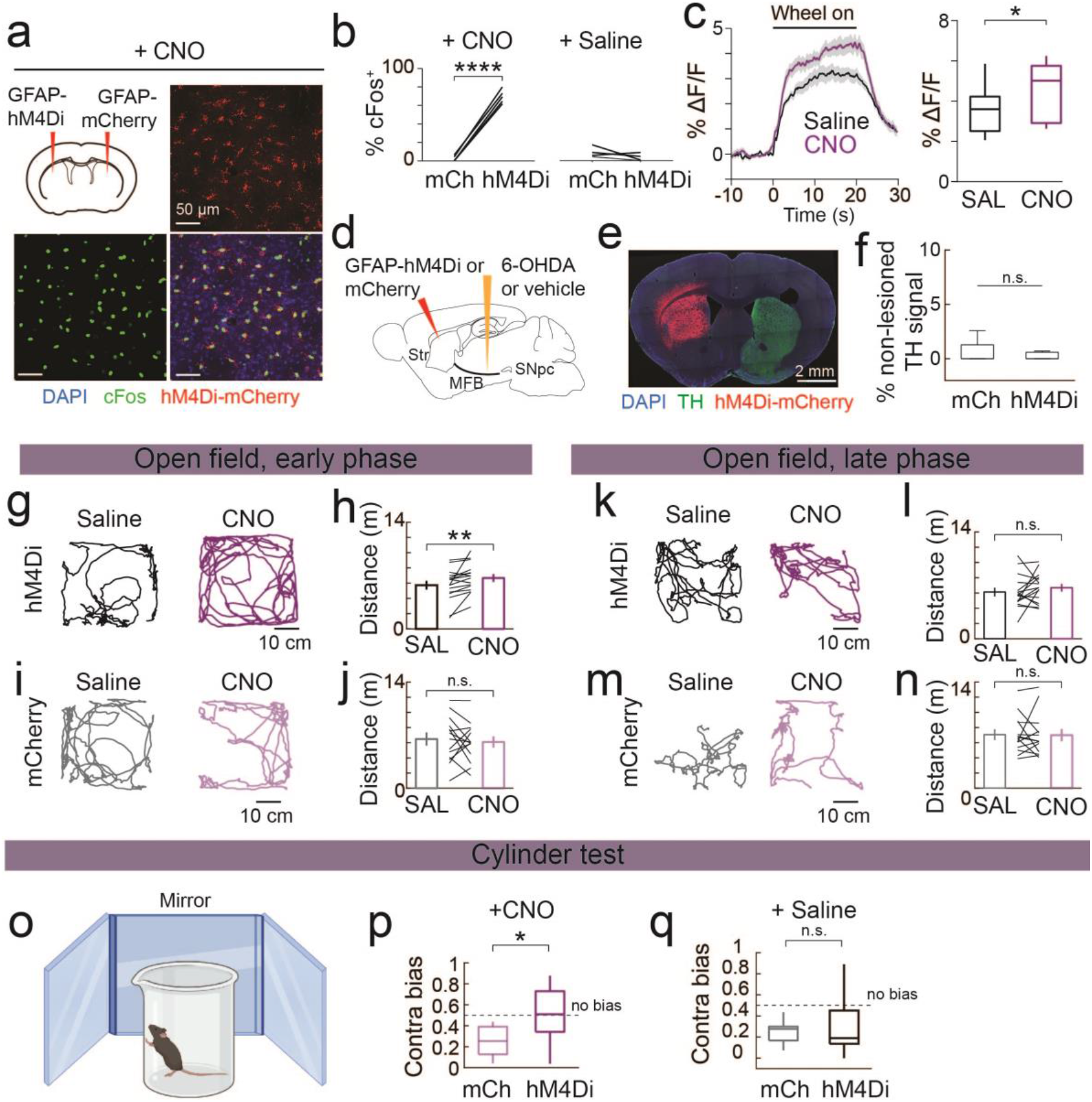
Chemogenetic activation of striatal astrocytes improves motor deficits from hemi-parkinsonian 6-OHDA lesioning. (**A**) Experimental schematic and representative histology showing c-Fos activation in hM4Di-expressing cells in the dorsal striatum following CNO administration (**B**) Percent of hM4Di and mCherry expressing striatal astrocytes showing immunoreactivity for c-Fos 2 hours after intraperitoneal injection of saline or CNO. (**C**) Astrocyte activity was recorded during locomotion on a motorized wheel in Aldh1l1+/lck-GCaMP6f+ transgenic animals injected with viral gfap-hM4Di-mCherry in the dorsal striatum. Comparison of astrocyte responses following saline (SAL) and CNO administration is shown. (n = 15 trials from 3 mice, *p* = 0.031017, *z* = -2.1569; Wilcoxon ranked-sum test) (**D**) Experimental schematic showing strategy to assay the effects of chemogenetic activation of striatal astrocytes in animals lesioned with 6-OHDA injection in MFB. Note that striatal virus injection was in the same cerebral hemisphere as the dopamine lesion. (**E**) Representative histology showing loss of tyrosine hydroxylase (TH) immunoreactivity and presence of hM4Di in animals for behavioral experiments. (**F**) Striatal TH immunofluorescence on the lesioned side normalized to the non-lesioned side for hM4Di and mCherry expressing mice (n = 13 for mCherry, 15 for hM4Di, *p* = 0.788, *z* = 0.269; Wilcoxon ranked-sum test). (**G**) Example locomotion trajectories during first 2 minutes (early phase) of the open field assay after saline and CNO injections from a 6-OHDA lesioned animal expressing astrocytic hM4Di. (**H**) Distance traveled in early phase for saline and CNO administration (n = 15 mice, *p* = 0.008, *z* = -2.67; Wilcoxon signed-rank test). (**I**, **J**) Same as **G** and **H**, but for mCherry expressing astrocytes (n = 13 mice, *p* = 0.60, *z* = 0.524). (**K**, **L**) Same as **G** and **H**, but for the late phase of the open field (mins 8-10; n = 15 mice, *p* = 0.46, *z* = -0.738; Wilcoxon signed-rank test). (**M**, **N**) Same as **K** and **L**, except for late phase of the assay (n = 13 mice, *p* = 0.806, *z* = 0.245; Wilcoxon signed-rank test). (**O**) Schematic of cylinder task to assess contralateral paw usage in dopamine lesioned animals. Mice were recorded with mirrors surrounding the cylinder to aid in the determination of initial paw usage. (**P**) Proportion of rears using the paw contralateral to the side of 6-OHDA lesion (contra bias) in either hM4Di or mCherry expressing animals after CNO injection (n = 13 hM4Di and 11 mCherry expressing mice; *p* = 0.015, *z* = 2.43; Wilcoxon ranked-sum test). (**Q**) Contra bias of mCherry and hM4Di expressing mice after saline administration (n = 11 hM4Di and 12 mCherry mice, *p* = 0.975, *z* = -0.031; Wilcoxon ranked-sum test).

We further tested how chemogenetic activation affects locomotion responses of striatal astrocytes. We virally expressed GFAP-hM4Di-mCherry in transgenic mice stably expressing GCaMP6f-lck in astrocytes to avoid potential confounds with expressing multiple viruses. After training on the motorized wheel, we used fiber photometry to measure locomotion related astrocyte responses 30 mins after treatment with saline or CNO. Chemogenetic activation increased locomotion-related astrocyte activity (**Fig. 4c**). Hence, chemogenetics facilitates astrocyte calcium activity.

### Functional astrocyte activation improves motor deficits in hemi-parkinsonian mice

Our findings thus far raise the question of whether the astrocyte activity deficits observed with dopamine loss contribute to movement deficits in Parkinson’s disease. Given the facilitatory effect of chemogenetic activation on astrocyte activity (**Figs. 4a-c**), we tested how this manipulation impacts behavior in hemi-parkinsonian and sham mice. Subgroups of mice receiving either vehicle (sham) or 6-OHDA (lesioned) in the MFB were injected with AAV viruses expressing GFAP-hM4Di-mCherry or AAV-GFAP-mCherry in the dorsal striatum (Figs 4d and 4e). 4-5 weeks after surgery, we gave mice I.P injections of saline or CNO 30 minutes prior to performing the open field test for 10 mins. We first determined how dopamine loss itself affects open field behavior by comparing mCherry expressing sham and lesioned mice in the saline treatment condition. As expected, dopamine loss reduced the total distance traveled (**Figs. S4a** and **S4b**). Open field locomotion partly reflects exploration of a novel environment and subsides with habituation to the arena over time^43^. Lesioned animals showed a higher decrease in distance traveled during early (first 2 minutes) compared to the late phase of the open field assay (last 2 minutes; **Figs. S4c** and **S4d**). Hence, dopamine loss especially perturbs explorative locomotion.

Given the time-dependent effect of dopamine loss on open field performance, we tested how chemogenetic astrocyte activation affects locomotion during early and late phases of the assay. There was no difference in locomotion between saline and CNO conditions for DREADD expressing sham mice (**Figs. S5a-c**), presumably due to a ceiling effect of astrocyte recruitment in dopamine intact animals. In contrast, CNO treatment in DREADD expressing lesioned mice increased the distance traveled in the early phase of the open field assay compared to saline (Figs. 4g and 4h). No such modulation was observed in mCherry expressing lesioned or sham mice (Figs. 4i and 4j). We did not detect any difference between CNO and saline conditions in the late phase of the open field assay (**Figs. 4k-n**). Together, these results show that chemogenetic astrocyte activation improves open field exploration in dopamine lesioned mice.

We further evaluated the behavioral effects of functional astrocyte activation using the cylinder test (**Fig. 4o**), a widely used behavioral assay of motor asymmetry in hemi-parkinsonian rodent models^44^. Chemogenetic astrocyte activation significantly improved usage of the affected paw contralateral to the lesion side. While cylinder test performance was similar for saline injections between hM4Di and mCherry expressing mice, CNO treated hM4Di mice showed more contralateral movements than CNO treated mCherry mice (**Figs. 4p** and **4q**). We did not observe differences in cylinder test performance with sham mice (**Fig. S5d** and **S5e**). Hence, chemogenetic astrocyte activation improves motor asymmetry in hemi-parkinsonian mice. Taken together with our open field data, these results suggest that chemogenetic activation of striatal astrocytes improves behavioral deficits resulting from dopamine loss.

## Discussion

We show that striatal astrocytes are robustly activated during locomotion (**Figs. 1, 2**), that dopamine loss reduces locomotion-related astrocyte activity (**Fig. 3**), and functional astrocyte activation with chemogenetics improves motor deficits in dopamine lesioned mice (**Fig. 4**). The observed motor improvements with astrocyte manipulations in dopamine lesioned animals suggest that astrocytes represent a complementary pathway of neuromodulation to dopamine signaling. Together, our experiments provide a new framework for understanding striatum’s role in regulating PD-relevant motor behavior.

Dopamine lesioned animals show reduced voluntary locomotion (**Fig. S4**). Hence, we used a motorized wheel to compare sham and dopamine lesioned animals under similar locomotion conditions. Astrocytes responded largely similarly during spontaneous and forced locomotion in intact animals, reaching a steady state level of activation after a rising period that persisted throughout the locomotion bout. Under both conditions, response activation was faster with locomotion onset compared to deactivation with locomotion offset (**Fig. 1f**, **2g** and **2h**). While there was no modulation of astrocyte activity with running speed under either condition, responses were slightly modulated by bout duration in the forced locomotion condition but not during spontaneous locomotion (**Figs. 1j**, **1k**, and **3g**). These results are not unique to striatal astrocytes as locomotion produces calcium activity with similar response properties in cerebellar Bergmann glia and cortical astrocytes^37,45^.

Bulk locomotion-related calcium responses recorded with fiber photometry are slower in astrocytes compared to neurons (**Fig. 2g** and 2h). These results are consistent with previous studies comparing the time course of astrocyte and neuronal calcium activity at single cell resolution^24^ and suggest that these cell types may regulate behavioral processes at different timescales. One interpretation is that astrocyte activity signals a locomotion behavioral state for the striatal circuit and promotes continued coordination of incoming cortical and thalamic sensorimotor information^24,37,46^ with dopaminergic neuromodulation and local neuronal processing^47^. Reduction of astrocyte calcium activity with dopamine loss would disrupt this process, which in parallel with the well-documented effects of dopamine loss on neuronal intrinsic properties and synaptic structure and function^3^ would result in motor dysfunction.

We show a reliable reduction in locomotion-related astrocyte activity with dopamine depletion (**Fig. 3e-h**). Our current data does not distinguish whether these results are due to loss of direct action of dopamine on striatal astrocytes or indirectly through known changes in local neuronal function and synaptic inputs^48–50^. Striatal astrocytes express dopamine receptors in their distal processes, potentially within the range of neuronal synapses^2–4^. Moreover, dopamine activates cortical astrocytes via noradrenergic receptors^48^, which provides another avenue for dopaminergic modulation of astrocyte activity. On the other hand, we only detected a moderate decrease in astrocyte locomotion responses with acute pharmacological blockade of dopamine receptors in intact animals and L-DOPA had a consistent but limited effect on astrocyte activity in dopamine depleted animals. These results point to the possibility that astrocyte signaling works in parallel rather than downstream to dopaminergic neuromodulation. More work is required to understand the exact relationship between dopamine and astrocyte signaling and how their synergistic activity shapes striatal neuronal activity and function.

Locomotion in the open field is driven partly by exploration and subsides as animals habituate to the novel environment^49^. In agreement with previous work demonstrating that dopamine signaling is important for exploratory behavior^51^, we found that lesioned animals showed a pronounced deficit in the early phase of the open field when the exploratory drive is the highest (**Fig. S4d**). A recent study^52^ reported a similar deficit in open field exploratory behavior with intrastriatal 6-OHDA injections that produce milder dopamine loss than the MFB injections employed in our study, suggesting that disrupted exploratory behavior is a sensitive metric for striatal dysfunction following dopamine loss. We found that chemogenetic astrocyte activation in lesioned animals increased exploratory locomotion while the same manipulation had no effect on dopamine intact animals, possibly because astrocyte activity was maximally engaged in sham animals during open field exploration. One interpretation of this finding is that dopamine and astrocyte signaling complementarily contributes to exploratory locomotion.

Chemogenetic astrocyte activation also improved cylinder task performance (**Fig. 4o-p**), which is a well-validated behavioral assay for motor dysfunction in hemi-parkinsonian rodent models^45^. These results are particularly relevant to cases of asymmetrical PD symptomology seen in some clinical cases^51^, potentially revealing astrocytes as a therapeutic cell type for improvement of bilateral coordination in cases of excess DA neuron loss on one side of the brain. The striatum is critical for the coordination of bilateral movements and DLS specific lesions impair bilateral movement duration and trajectory^53–55^. Unilateral lesioning with 6-OHDA imbalances striatal output activity across the two cerebral hemispheres, leading to suppression of the lesioned side compared to the non-lesioned striatum^56^. Our astrocyte fiber photometry results in 6-OHDA lesioned animals mirror the broad reduction in ipsilateral striatal neuronal activity seen in previous work^57^. That our data show an improvement with asymmetrical paw usage with chemogenetic astrocyte activation suggests that astrocytes may restore some ipsilateral striatal output. Since exogenous activation of astrocytes in the lesioned striatum produces improvement in bilateral paw usage, our results further suggest that while astrocyte activity post-lesion is reduced, the cells still retain the ability to influence behavioral output when artificially stimulated via GPCR signaling. Relatedly, we found that L-DOPA administration increased locomotion-related astrocyte activity 4-5 weeks post dopamine lesioning during the 20s locomotion bout condition. This demonstrates that the reduction in activity is not permanent and can be at least partially restored. These findings further support the case for astrocytes as potential therapeutic targets given that the observed behavioral improvements with chemogenetics were without direct dopamine manipulations (i.e. dopamine neurons are still lost and dopamine transmission in the striatum was not directly manipulated). As exogenous DA infusion in the form of L-DOPA carries significant side effects, this complementary pathway to improving unilateral PD symptoms may carry clinical promise.

Overall, our data provide evidence for the role of striatal astrocytes as major components in shaping striatum-related behaviors such as locomotion and bilateral motor coordination after dopamine loss. Unlike DA neurons which are permanently lost with both 6-OHDA lesioning and in clinical PD, the astrocytic network is largely persevered. Although DA loss may switch astrocytes to an inflammatory reactive state^58,59^, exogenous activation of these cells via GPCR signaling still improves motor function (**Fig. 4**). Taken together, our experiments show that astrocytes represent a complementary pathway for targeting the motor symptoms of PD. Further work is necessary to develop new methods for targeting astrocyte calcium activity with higher precision.

## Acknowledgements

This work was supported by grants from the National Institute of Mental Health (R00MH112855 to R.H.), Parkinson’s Foundation (R.H.), American Parkinson Disease Association (R.H.), Brain Research Foundation (R.H.), and the National Institute on Alcohol Abuse and Alcoholism (T32AA028254 to W.E.). We thank Dr. Grayson Sipe, Dr. Cherish Ardinger, and Nithik Chintalacheruvu for helpful comments on this manuscript and Cielo Tumbokon for help with quantification of c-Fos cell counts. Biorender was used for creating figure schematics.

## Competing interests

The authors report no competing interests.

## Author contributions

W. Evans, S. Baskar, and R. Huda designed experiments. W. Evans performed and analyzed the fiber photometry and histology experiments. S. Baskar performed and analyzed the behavioral and histology experiments. R. Huda analyzed the fiber photometry and behavioral experiments. A.R. Castro E Costa, A. Arigbe, and S. Ravoori aided with histology, fiber photometry, and behavioral experiments. W. Evans and R. Huda wrote the manuscript with edits from all authors. R. Huda supervised the study.

## Data and code availability

All data and code generated as part of this study are available from the corresponding author upon reasonable request.

## Methods

All experimental procedures performed on animals were approved by the Rutgers University Institutional Animal Care and Use Committee (Protocol #202000004). C57BL/6J mice were purchased from the Jackson Laboratory at 7-9 weeks old, and additional mice were bred in-house. Animals were acclimated to the housing facility for at least one week before performing stereotaxic surgeries. Following surgeries, mice were housed in a reversed 12-hour light/dark cycle in standard sized cages and provided food and water ad libitum.

### General stereotaxic surgery procedures

Procedures were as described previously^59^. Briefly, mice were deeply anesthetized under 4% isoflurane and then mounted on a heated stereotaxic frame (∼37°C, Stoelting). Anesthesia was maintained with 1–2% isoflurane and adjusted as needed. Mice were given extended-release buprenorphine (Ethiqa-XR, 3.25mg/kg) to provide analgesia for up to 72 hours; meloxicam (10mg/kg) was provided if additional analgesia was required during surgical recovery. Bupivacaine (1 mg/kg) was given subcutaneously under the scalp. Scalp hair was removed with depilatory cream (Nair), and surgical areas were disinfected with 3x alternating application of betadine and 70% ethanol. A midline incision was made, and the skull was leveled relative to bregma and lambda in the dorsoventral and mediolateral axes. Mice for behavioral experiments were group housed; animals made for fiber photometry recordings were singly housed. Mice recovered for at least one week post-surgery before starting experiments.

### 6-OHDA infusion

Mice were dosed with desipramine hydrochloride (HCl) and pargyline HCl (10ml/kg) via intraperitoneal injection thirty minutes prior to 6-OHDA infusion^60^. A burr hole was drilled in the skull, and 1000 nL of 6-OHDA HBr (3.6 mg/ml), mixed in a vehicle solution of 0.02% ascorbic acid (pH 7.4) in 0.9% sterile saline, was pressure injected (Nanoject III, Drummond) into the medial forebrain bundle through the opening made at the following coordinates defined relative to bregma: AP: -1.2, ML: +1.2, DV: -5.0 (from bregma). Sham animals received both the intraperitoneal injection of the drug cocktail and an ascorbic acid intracranial injection. Injections were made over 10 min, and the needle was left in place for another 10 min. The incision was sutured shut and mice were kept warm while they recovered from anesthesia. Mice were monitored for 14 days post surgery. They received daily subcutaneous injections of 5% dextrose in saline and DietGel Boost mixed with powdered chow in their home cage to aid in healthy recovery.

### AAV injections

A burr hole was made in the skull over the target site using a handheld dental drill. Viruses were pressure injected with a Nanoject III (Drummond) through a glass micropipette slowly lowered into the craniotomy over the course of 5 minutes. For behavior experiments, 500nL of either AAV5-GFAP-hM4D(Gi)-mCherry (Addgene catalog # 50479-AAV5) or AAV5-GFAP-mCherry (Addgene # 58909-AAV5) was injected into the left dorsal striatum (coordinates: AP: +1.0mm, ML: +1.7mm, DV: -2.6mm, relative to bregma). For fiber photometry experiments, 500nL of either AAV1-syn-GCaMP8m (Addgene # 162375-AAV1) or AAV5-gfaABC_1_D-lck-GCaMP6f (Addgene # 52924-AAV5) was injected into the left dorsal striatum at either AP: 1.0, ML: 1.5, DV: -2.7 (dorsomedial striatum, DMS) or AP: 0.4, ML: 2.3, DV: -2.7 (dorsolateral striatum, DLS). After injection the pipette was kept in place for an additional 5 minutes before removing slowly. Incisions were sutured or tissue glued (Vetbond, 3M) and mice were kept warm while they recovered from anesthesia. Animals were monitored for at least 3 days post procedure to ensure healthy recovery.

### Fiber optic implants

After AAV injection into the dorsal striatum was complete, a fiber optic cannula (Ø1.25mm ceramic ferrule, 400µm Core, 0.39NA, RWD Life Sciences) was fixed onto a stereotaxic cannula holder (ThorLabs) and slowly lowered into the same striatal craniotomy as the viral injection. It was implanted 0.1mm above the AAV injection in the dorsoventral axis, and dental cement (MetaBond, Parkell) mixed with black ink was applied to secure the fiber optic in place. A custom headplate (eMachineShop) was attached to the skull using cyanoacrylate (Krazy Glue) and dental cement (MetaBond, Parkell) to enable head-fixation during experiments. Sterile saline was administered subcutaneously, and the mice were kept warm while they recovered from anesthesia. Animals were monitored for at least 3 days post procedure to ensure healthy recovery.

### Fiber photometry

Animals surgically prepared for fiber photometry recordings were habituated to both our voluntary and forced locomotion apparatuses for at least 3 practice sessions before data collection. Striatal astrocytes or neurons were transfected with a Ca^2+^ indicator (syn-GCaMP8m or gfaABC_1_D-lck-GCaMP6f) and allowed at least 3 weeks for expression. GCaMP fluorophores were excited with both a 465 nm and 405 nm LED through the same patch cord and emission was recorded using a Tucker-Davis Technologies RZ10X system. Fluorescent emission was separated into Ca^2+^ dependent and isosbestic signals using a Doric cube. During recordings the patch cord was tightly adhered to the implanted ceramic ferrule using a ceramic coupling (Doric Lenses) to accurately read incoming signal. Photometry signals were processed in a custom Python script and ΔF/F was calculated using the least-squares polynomial fit of the isosbestic channel to the signal channel. Forced locomotion experiments were analyzed in Python by averaging together trials of the same duration per animal and metrics were then calculated from the averaged trace.

### Head-fixed voluntary locomotion

We built a behavioral rig for voluntary head-fixed locomotion experiments. A plastic running wheel was mounted to the shaft of a rotary encoder (Yumo) using a custom designed 3D printed adaptor. The rotary encoder was housed in a 3D printed enclosure and mounted on an optical breadboard (Thorlabs). The floor of the enclosure had a 20° angle to slightly tilt the running wheel. Head-plate holders were mounted so that the forepaws of head-fixed mice were on upward part of the slope. We found that this design promoted robust voluntary running.

Rotary encoder output was connected to an Arduino UNO running a custom sketch. The rotary encoder was queried at 100 Hz and wheel movements were recorded on a PC computer running MATLAB scripts. Wheel movement recordings and fiber photometry signals were synchronized using the same MATLAB script by initiating wheel recording on the Arduino at the same time as a TTL signal was sent to the DAQ board of the fiber photometry system. Animals were habituated to voluntary wheel running for at least 3 sessions until they consistently ran forward; animals which did not become proficient in forward running were excluded from analysis.

### Head-Fixed Forced Locomotion

A circular treadmill was attached to a D/C rotary motor and controlled with a custom-built circuit and MATLAB script. Locomotion bouts were spaced one minute apart and were randomized among 2, 5, 10, and 20s for 5 bouts per session or for 10s 10 times for direct comparison of cell type and regional differences (**Fig. 2**). Locomotion bouts were aligned post-hoc using TTL signals fed into Synapse software (TDT) from the wheel control circuit. Animals were habituated to running on the motorized wheel for 3 days before beginning experimental recording sessions. For experiments involving pharmacology, agents were injected 30-40 minutes prior to experiments.

### Open Field Test

The open field test was performed in a 40 x 40cm arena (60101, Stoelting) on days 25 and 35 post 6-OHDA lesion surgery. Thirty minutes before the first test mice were given IP injections of either CNO or saline and received the opposite injection before the second test. Mice were then placed in the center of the open field arena and exploratory behavior was video recorded for 10 minutes at 25 fps. A DeepLabCut^61^ model was trained to track five keypoints on the mouse: nose, right ear, left ear, body center, and base of the tail. Distance traveled was computed using the body center keypoint. Four corners of the open field box were also tracked. These points were used to calculate the length of each side of the box in pixels. X-Y pixel coordinates of the body center keypoint were converted to cm using the known cm length of each box side (40cm). Points tracked with a <95% probability were eliminated, and the remainder of the data interpolated using the ‘interp1’ MATLAB function. X-Y coordinates were LOESS filtered using the function ‘smooth’ with a window size of 10 frames. Coordinates were then used to calculate the frame-by-frame distance, which were summed to calculate the total distance traveled across various time bins according to the reported analyses.

### Cylinder Test

The cylinder test was performed on days 21 and 28 days after the 6-OHDA lesion surgery. Thirty minutes before the first test, mice received intraperitoneal injections of either saline or Clozapine N-oxide (CNO), and the opposite injection prior to the second test. The mouse was placed into a 6.5 x 4in glass cylinder with three mirrors facing an infrared camera to enable 360-degree view of rearing behavior which was recorded in the cylinder for 20 minutes. Videos were manually scored using ImageJ. Each rearing event was characterized by these criteria: Bilateral or Unilateral, and First Paw (Right or Left). Unilateral events were recorded when a mouse reared up on its hind legs and placed one paw on the glass during a weight-bearing rear, whereas bilateral events were recorded when the mouse used both paws for the rear. Further characterization of the ipsi-vs contra-paw usage during a unilateral event was recorded based on which paw was on the glass during the weight-bearing portion of the rear.

### Immunohistochemistry

Mice were perfused with saline and 4% paraformaldehyde (PFA). The extracted brains were stored in PFA for 24h additional fixation. The brains were sectioned using a compresstome (VF-310-0Z, Precisionary Instruments), and 60μm thick coronal slices of the dorsal striatum and substantia nigra were collected and stored in 12-well plates containing PBS. Slices were washed in PBS thrice for five minutes each and then blocked in 3% normal goat serum in PBST-Az solution for 2 hours. Following this, chicken anti-TH 1:1000 in PBST-Az (TYH-0020, Aves Lab), rabbit anti-c-Fos 1:1000 in PBST-Az (226008, Synaptic Systems), rabbit anti-NeuN 1: 500 (Synaptic Systems), or mouse anti-S100b 1:1000 (Synaptic Systems) was placed in the blocking solution and left overnight at room temperature on an orbital shaker. Slices were washed in PBS thrice for five minutes each. Finally, slices were placed in a secondary antibody solution of Alexa Fluor 488 goat anti-chicken IgY 1:500 (A11039, Invitrogen), Alexa Fluor 647 goat anti-rabbit IgY 1:1000 (A32733, Invitrogen), Alexa Fluor 488 goat anti-rabbit IgY 1:500 (A11008, Invitrogen), or Alexa Fluor 647 goat anti-mouse IgY 1:500 (A21235, Invitrogen) for 2 (NeuN) or 4 (all other stains) hours on an orbital shaker at room temperature. Slices were mounted on permafrost slides with DAPI mounting medium and covered with 24 x 55mm No.1 cover glass and stored in a slide box in a 4°C refrigerator.

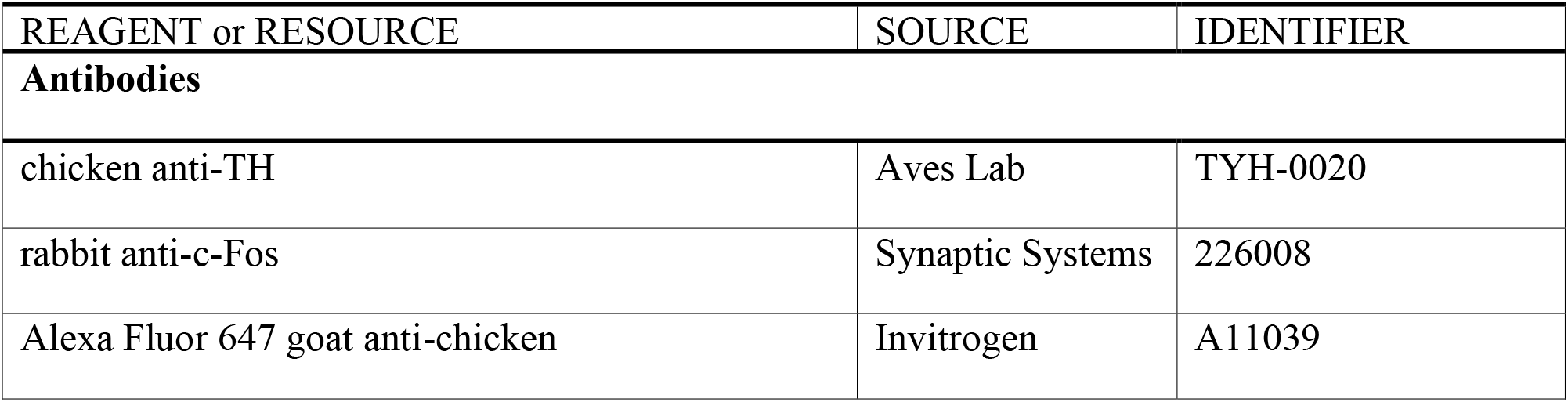
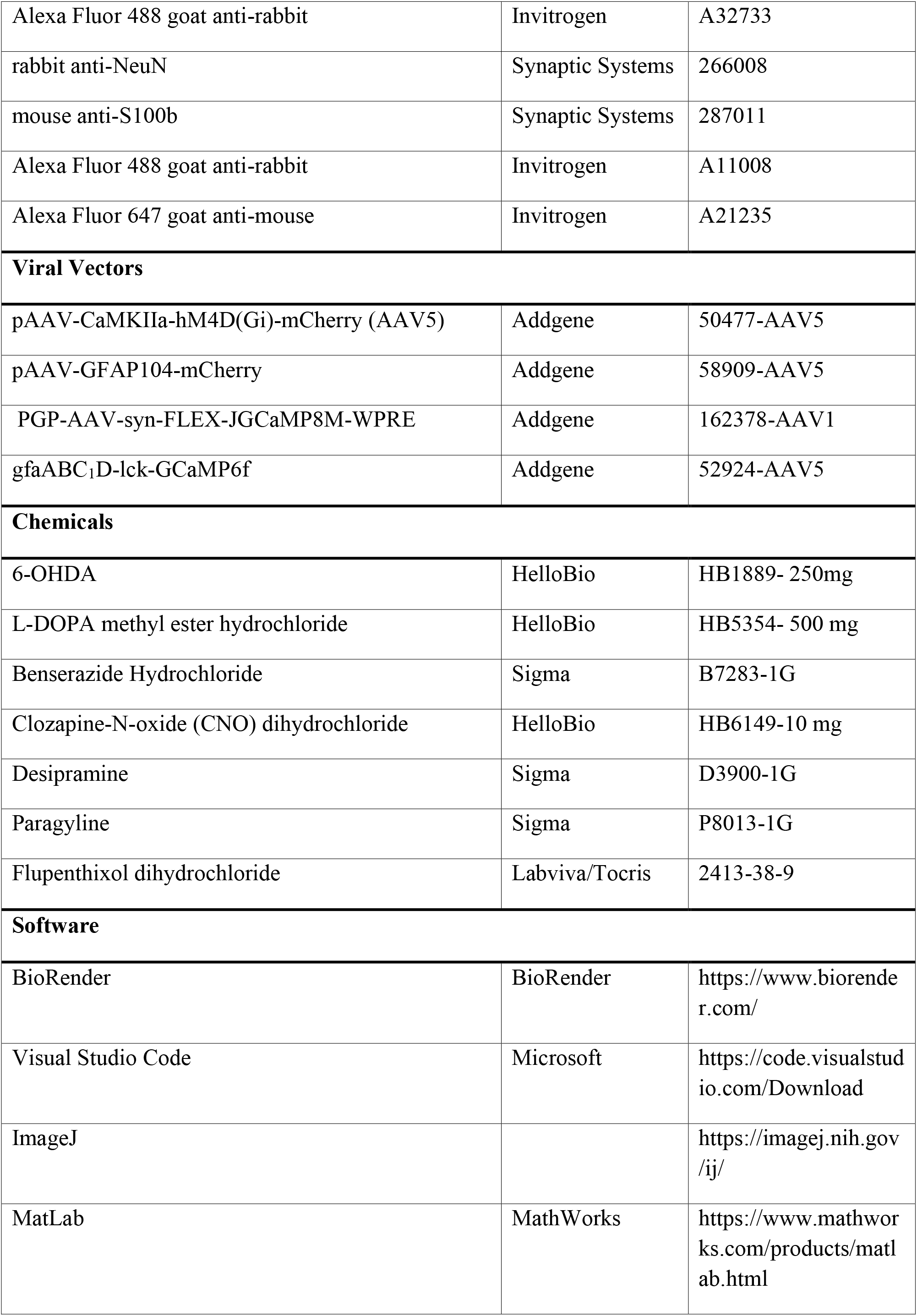
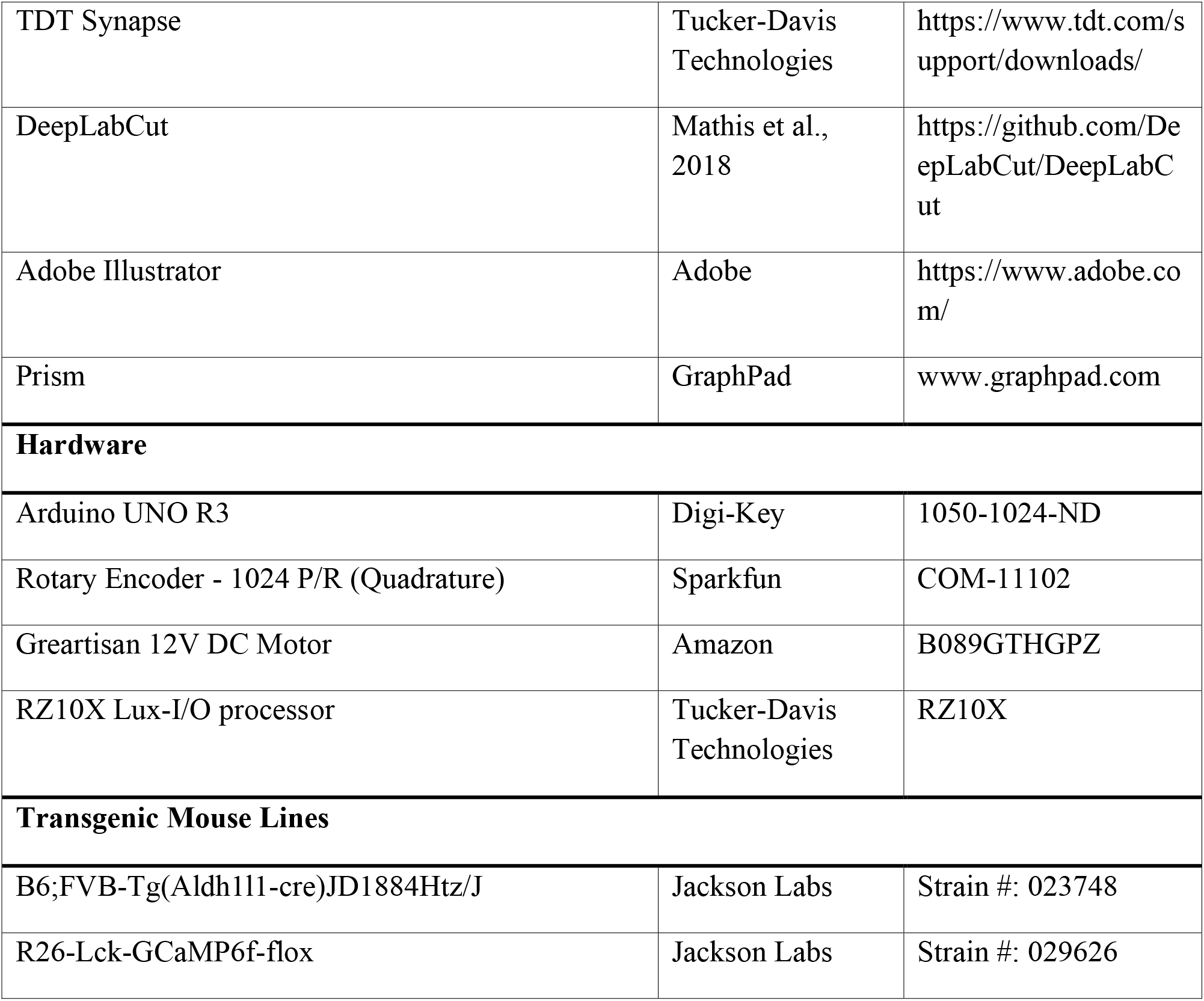
Key resources table

## Supplementary Figures

**Figure S1.**
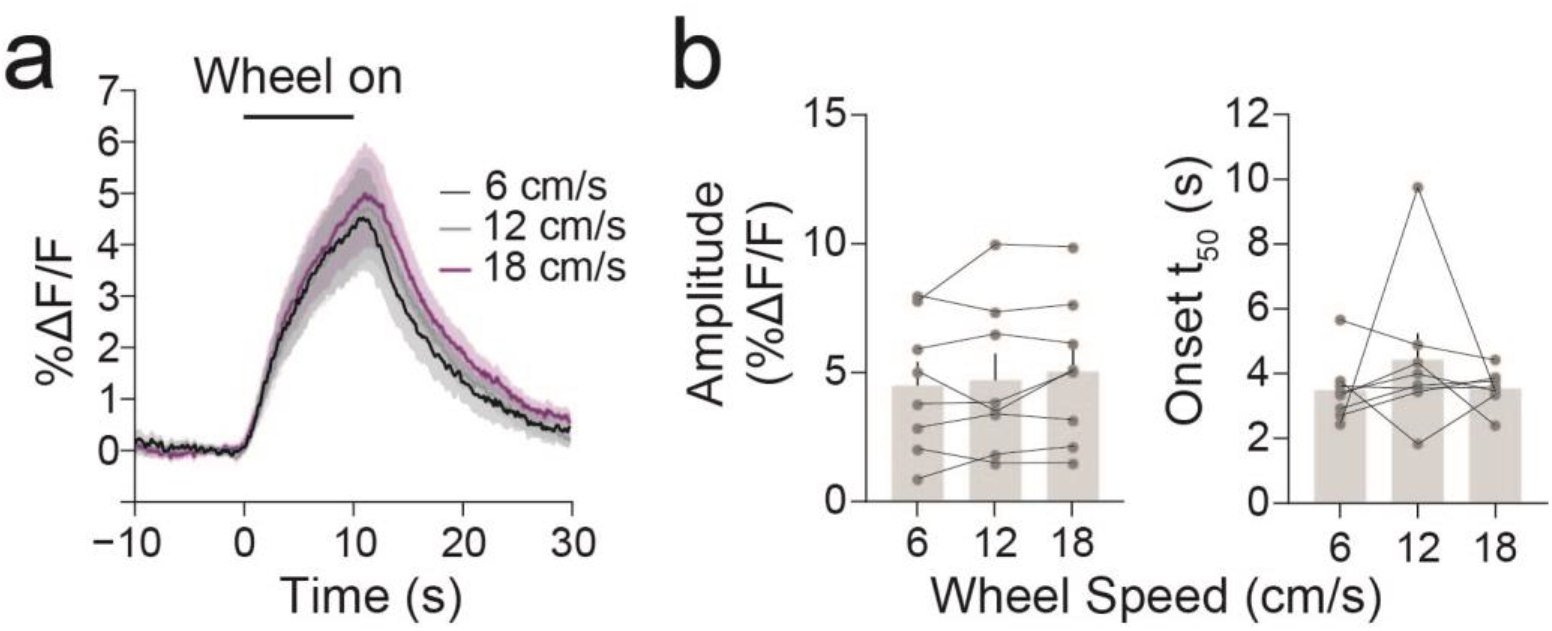
Motorized wheel speed does not modulate DLS astrocyte locomotion responses. (**A**) Trial-aligned locomotion responses at 6 cm/s (black), 12 cm/s (gray), and 18 cm/s (purple). (**B**) *Left*, peak locomotion response across wheel speeds (*F*(2, 15) = [1.353], *p* = 0.2903; repeated measures ANOVA). *Right,* time to 50% of peak response across velocities (*F*(2, 23) = [0.9505], *p* = 0.4101; repeated measures ANOVA).

**Figure S2.**
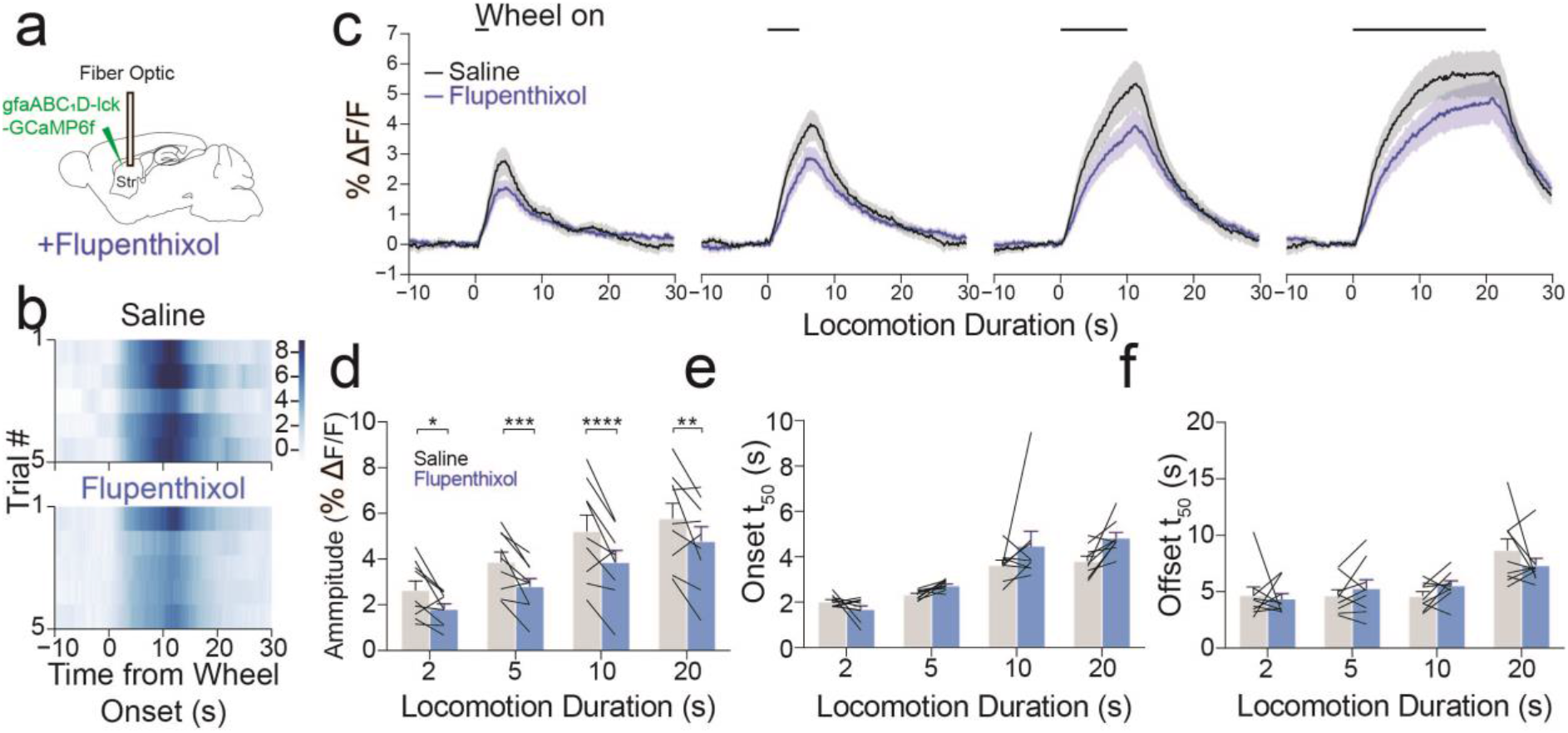
Pharmacological antagonism of dopamine receptors reduces locomotion-evoked astrocyte responses. (**A**) Experimental schematic showing injection of gfaABC_1_D-lck-GCaMP6f into dorsal striatum and placement of fiber optic for recording of fiber photometry signals. (**B**) Heatmaps showing single trial responses during 10s locomotion bouts from the same animal injected with either saline or flupenthixol. (**C**) Trial-averaged ΔF/F traces for all 4 locomotion durations with saline and flupenthixol I.P. injections. (**D**) Response amplitude was decreased with systemic flupenthixol administration (n = 9 mice, main effect of drug treatment *p* = 0.0064, interaction *F*(3, 24) = [0.781139], Sidak’s multiple comparison test 2s *p* = 0.0112, 5s *p* = 0.0010, 10s *p* < 0.0001, 20s *p* = 0.0021; two-way repeated measures ANOVA). (**E**) Time to 50% of peak ΔF/F was unaffected by flupenthixol administration (n = 9 mice, main effect of treatment *F*(1, 8) = [13.3654], p = 0.068; interaction *F*(3, 24) = 2.738, *p* = 0.0656; two-way repeated measures ANOVA). (**F**) Time to 50% offset from peak ΔF/F was unchanged with flupenthixol injection (n = 9 mice, main effect of treatment *F*(1, 8) = 0.0027, *p* = P=0.96; interaction *F*(3, 24) = 1.706, *p* = 0.1925; two-way repeated measures ANOVA).

**Figure S3.**
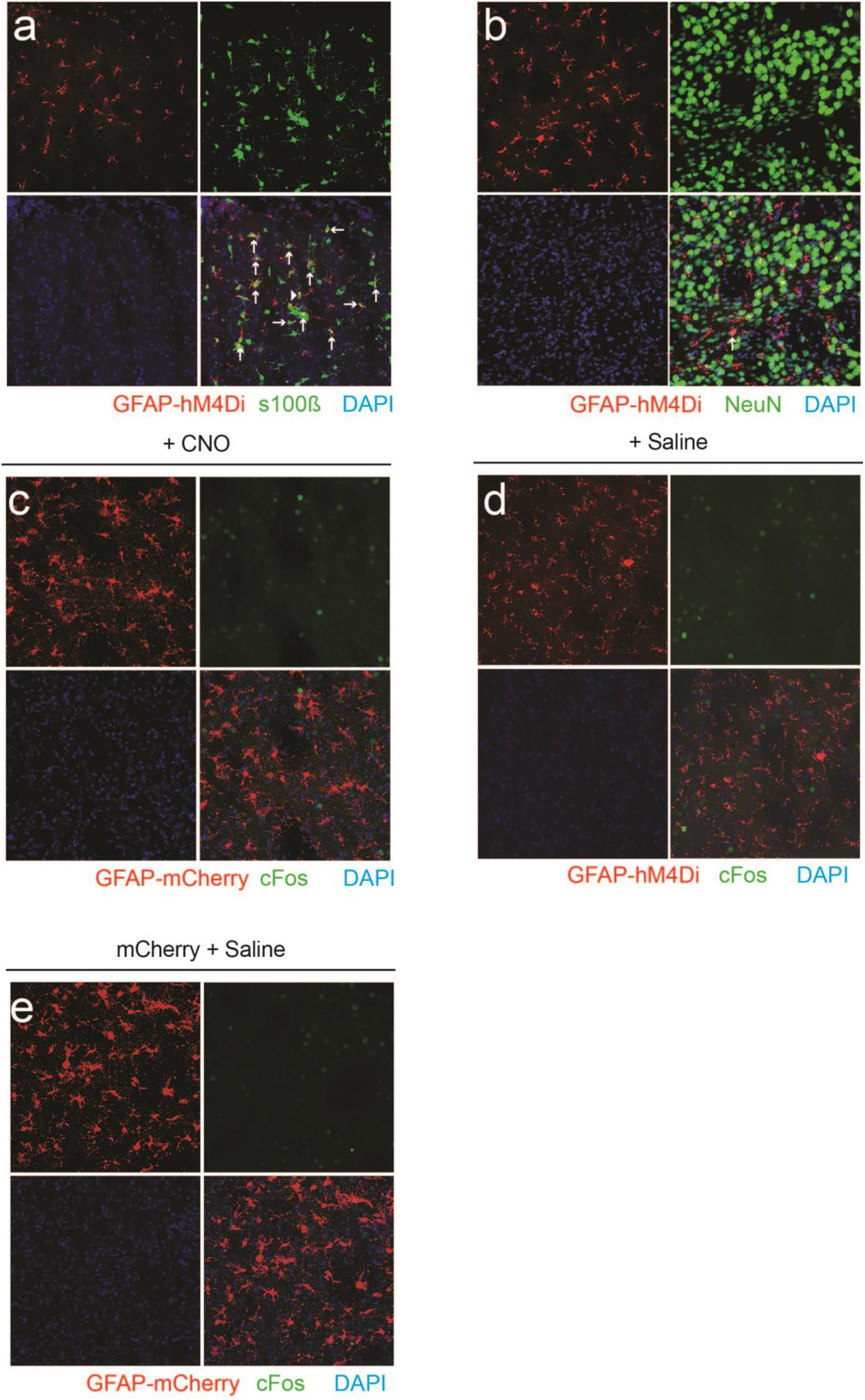
Representative histology from control groups for cFos immunoreactivity activation with hM4Di and CNO. (**A, B**) Overlap of viral GFAP-hM4Di-mCherry with the astrocyte marker S100β (**A**) and the neuronal marker NeuN (**B**). (**C**) The animal shown in **Fig. 4a** was injected with GFAP-mCherry in the opposite dorsal striatum. CNO administration did not produce cFos immustaining in mCherry labeled cells in this hemisphere. (**D, E**) Example images showing hM4Di-mCherry+ (**D**) and mCherry+ (**E**) striatal astrocytes and cFos immunostaining from animals I.P. injected with saline 2 hours prior to perfusion.

**Figure S4.**
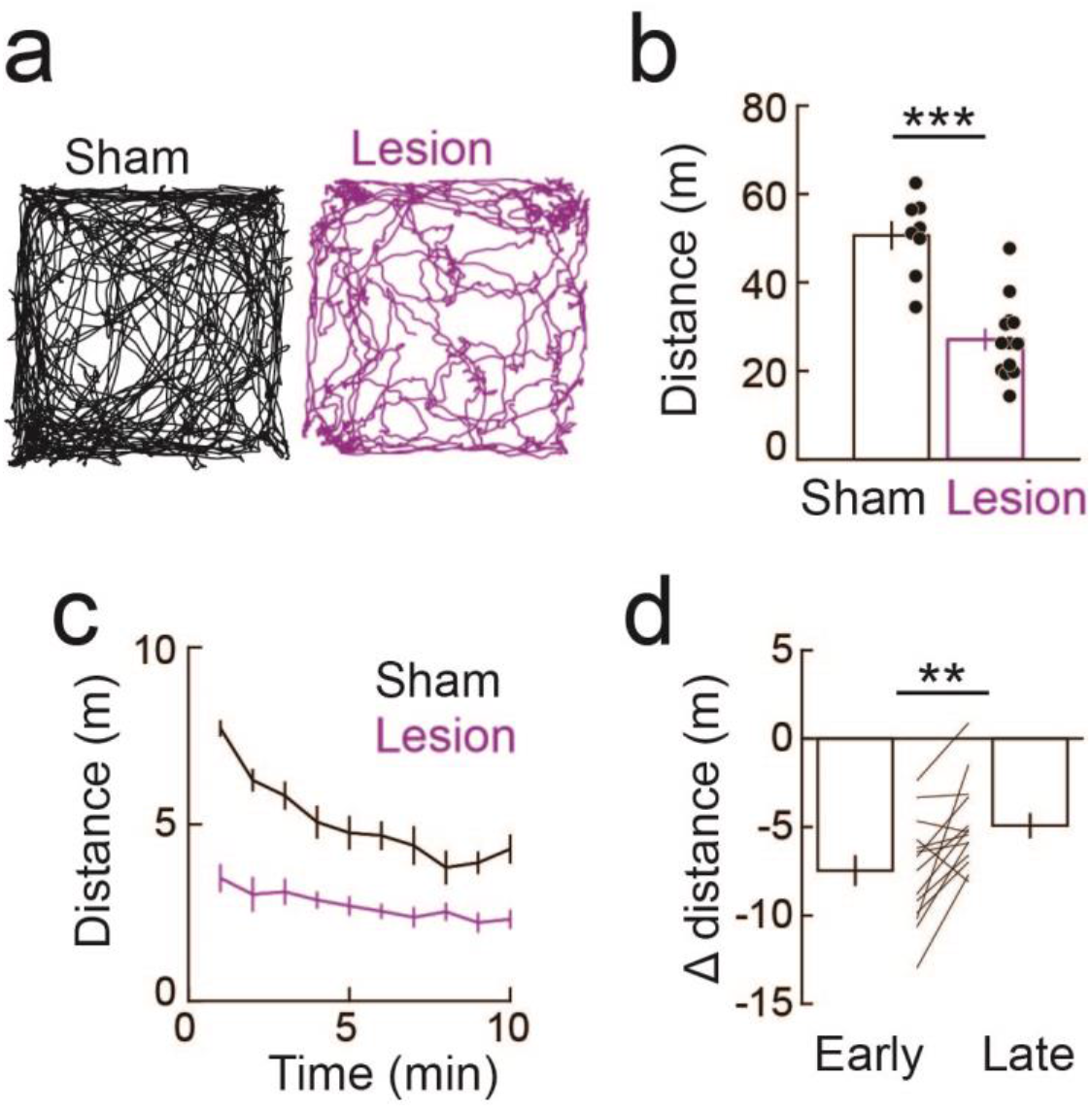
Time-dependent effect of dopamine loss on open field exploratory behavior. **(A)** Locomotion trajectories from example sham and dopamine lesioned mice. (**B**) Comparison of total distance traveled during the 10 min open field assay between sham and lesioned animals (n = 8 sham and 13 lesioned mice; \*\*\**p* = 0.0004, *z* = 3.51; Wilcoxon ranked-sum test). (**C**) Distance traveled in 1 min bins for sham and lesioned animals. (**D**) Delta distance, computed by subtracting the group averaged sham distance from individual lesioned animals, plotted for early (first 2 mins) and late (last 2 mins) phases of the open field assay (n = 13 mice, \*\**p* = 0.009, *z* = -2.62; Wilcoxon signed-rank test).

**Figure S5.**
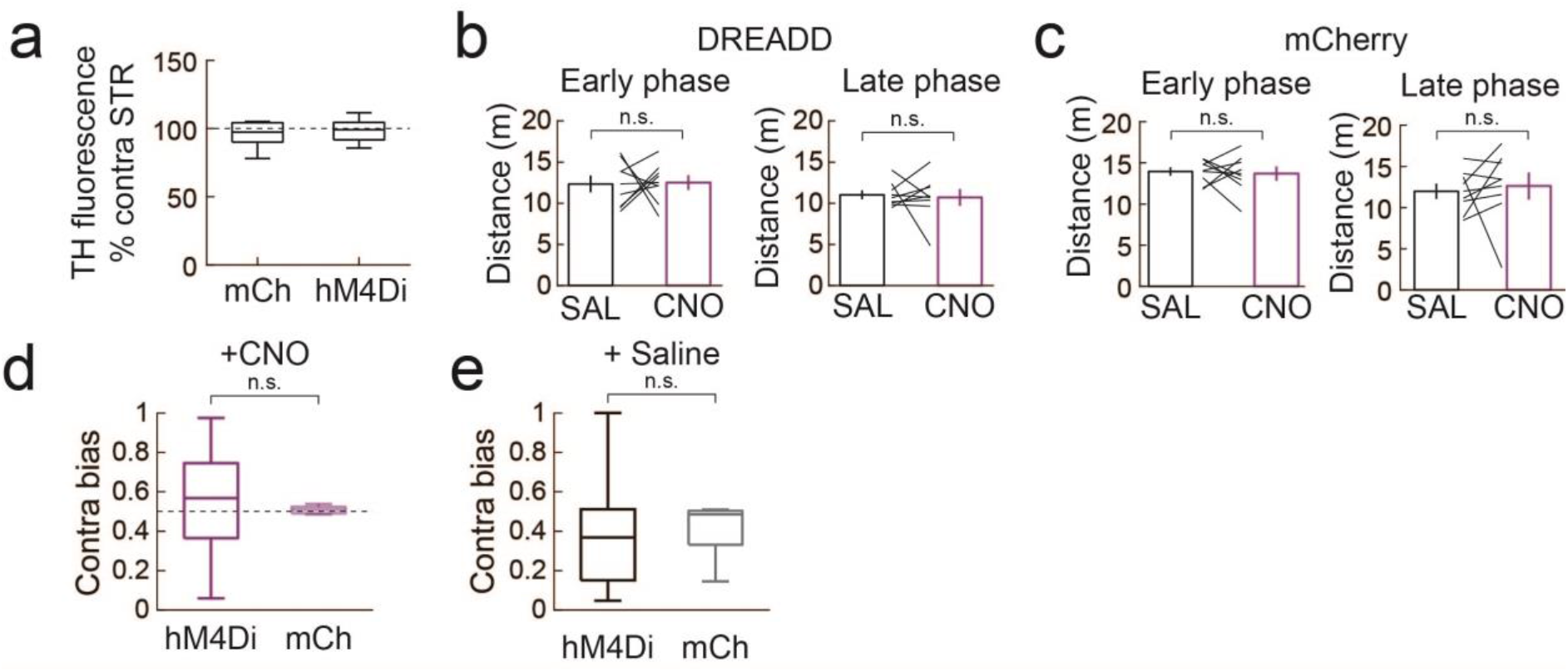
Effect of chemogenetic astrocyte activation on open field and cylinder test performance for sham treated mice. (**A**) TH immunofluorescence from mCherry (mCh) and hM4Di expressing mice. The immunofluorescence on the injected side was normalized by the non-injected side. (**B**) Distance traveled in the first 2 mins (early) and last 2 mins (late) of the open field assay following saline and CNO I.P. injections in hM4Di expressing sham mice (n = 8 mice; early: *p* = 0.889, *z* = -0.140; late: *p* = 0.779, *z* = -0.280; Wilcoxon signed-rank test). (**C**) Same as B, except for mCherry expressing mice (n = 8 mice; early: *p* = 0.889, *z* = 0.140; late: *p* = 0.327, *z* = -0.980; Wilcoxon signed-rank test). (**D**) Cylinder test performance with CNO adminstration in sham mice (n = 8 mice for both hM4Di and mCherry; *p* = 0.318, *z* = 0.998; Wilcoxon ranked-sum test). (**E**) Same as **D**, but for saline administration (n = 8 mice for both hM4Di and mCherry; *p* = 0.270, *z* = 1.10; Wilcoxon signed-rank test).

## Notes

### Competing Interest Statement

The authors have declared no competing interest.

